# Using pressure-driven flow systems to evaluate laser speckle contrast imaging

**DOI:** 10.1101/2022.09.16.508276

**Authors:** Colin T. Sullender, Adam Santorelli, Lisa M. Richards, Pawan K. Mannava, Christopher Smith, Andrew K. Dunn

## Abstract

**Significance:** Microfluidic flow phantom studies are commonly used for characterizing the performance of laser speckle contrast imaging (LSCI) instruments. The selection of the flow control system is critical for the reliable generation of flow during testing. The majority of recent LSCI studies using microfluidics used syringe pumps for flow control.

**Aim:** We quantified the uncertainty in flow generation for a syringe pump and a pressure-regulated flow system. We then assessed the performance of both LSCI and multi-exposure speckle imaging (MESI) using the pressure-regulated flow system across a range of flow speeds.

**Approach:** The syringe pump and pressure-regulated flow systems were evaluated during stepped flow profile experiments in a microfluidic device using an inline flow sensor. The uncertainty associated with each flow system was calculated and used to determine the reliability for instrument testing. The pressure-regulated flow system was then used to characterize the relative performance of LSCI and MESI during stepped flow profile experiments while using the inline flow sensor as reference.

**Results:** The pressure-regulated flow system produced much more stable and reproducible flow outputs compared to the syringe pump. The expanded uncertainty for the syringe pump was 8–20× higher than that of the pressure-regulated flow system across the tested flow speeds. Using the pressure-regulated flow system, MESI outperformed single-exposure LSCI at all flow speeds and closely mirrored the flow sensor measurements, with average errors of 4.6 ± 2.6% and 15.7 ± 4.6%, respectively.

**Conclusions:** Pressure-regulated flow systems should be used instead of syringe pumps when assessing the performance of flow measurement techniques with microfluidic studies. MESI offers more accurate relative flow measurements than traditional LSCI across a wide range of flow speeds.

## 1 Introduction

Laser speckle contrast imaging (LSCI) is an optical imaging technique based on coherent dynamic light scattering that has been widely adopted for visualizing changes in blood flow.^1, 2^ Requiring only a laser for illumination and a camera for detection, LSCI can create full-field maps of motion with high spatiotemporal resolution. This has produced great interest in using LSCI for imaging flow in preclinical neuroscience research^3–6^ as well as a broad set of clinical applications spanning the skin,^7, 8^ retina,^9, 10^ and brain.^11–13^ Efforts to improve the quantitative accuracy of LSCI resulted in the development of multi-exposure speckle imaging (MESI),^14^ which offers more robust estimates of flow compared to traditional single-exposure LSCI.^15, 16^ The acquisition of data at multiple camera exposure times increases the dynamic range of flow sensitivity while the more comprehensive dynamic light scattering model better accounts for the presence of static scatterers and instrumentation factors. These improvements have facilitated the chronic imaging of neurovascular blood flow^17–20^ and offer increased sensitivity during neurosurgical measurements.^21^

Due to its popularity, LSCI instrument development is an active area of research with microfluidic studies commonly used to characterize performance in controlled flow environments. Typically, studies designed to mimic blood flow involve the use of a flow channel embedded in a polydimethylsiloxane (PDMS) substrate,^14^ glass capillary tubing,^22^ or clear plastic tubing.^23^ More complex microfluidic networks have also been fabricated in glass, plastic, or epoxy substrates to represent superficial heterogenous vasculature.^24^ A scattering solution^14, 22^ or whole blood^23^ is then flowed through the channel at controlled rates and imaged using LSCI. For applications in rodent studies, 0.1 – 10 mm/s is a commonly selected range of flow speeds that corresponds with experimentally measured capillary speeds.^25^ This speed can be converted to a volumetric flow rate using the cross-sectional area of the flow channel, which is then used to program the flow control system.

Syringe pumps are the most widely used flow control system in microfluidic studies and are believed to provide a highly-controlled flow output. A survey of over twenty LSCI studies using microfluidics that were published since 2015 found that syringe pumps were used in the majority (90%) of experiments. The devices were from a variety of manufacturers and featured various syringe sizes and materials. Syringe pumps operate by applying mechanical force on the plunger of a syringe at a controlled speed. A rotating lead screw driven by a stepper motor controls the velocity of the plunger, which is calculated based on the syringe size and inner diameter to achieve a desired flow rate. However, syringe pumps produce oscillations in flow with varying frequency and magnitude due to the physical motion of the motor and from imperfections on the lead screw.^26, 27^ Syringe pumps have also been shown to have slow responsivity and can take a long time to achieve programmed flow rates.^28^ The trade-off between stability and responsivity makes it difficult^29^ to obtain both stable flow and a fast flow response with a syringe pump system, both of which are necessary for LSCI characterization measurements.

Pressure-driven flow control systems overcome the limitations of syringe pumps by eliminating the use of a motor and lead screw to provide very stable flow.^26, 28, 30^ These systems use pressure-regulated air to push liquid from a reservoir through the microfluidic device at a constant flow rate.^30^ In order to control the applied pressure, an absolute flow sensor must be added inline with the microfluidic device. Thermal mass flow sensors are commonly used to measure the low flow rates found in microfluidics. These sensors use heating and temperature-sensing elements to detect the magnitude and direction of flow and are factory-calibrated to measure absolute flow rates for various fluids. Integration of the flow sensor with a feedback loop allows for the programming of specific flow rates rather than pressure levels.

In this paper, we address two primary objectives. First, we quantify the uncertainty in flow generation for both a syringe pump and pressure-regulated flow system in microfluidic flow phantoms commonly used with optical blood flow imaging instruments. By understanding the real-world uncertainty associated with each flow controller, we can design a microfluidic study to characterize the performance of optical imaging techniques that minimizes the impact of flow-related errors. Secondly, we use the pressure-regulated flow system to assess the performance of both LSCI and MESI across a range of physiologically-relevant flow speeds. Using the inline flow sensor as reference, we demonstrate the superior accuracy of relative flow measurements with MESI compared to traditional single-exposure LSCI. Such measurements would have been confounded by the large uncertainties associated with syringe pump flow generation. The results of this study indicate that highly-accurate flow controllers, such as pressure-regulated flow systems, should be used when characterizing laser speckle techniques with microfluidic studies.

## 2 Methods

### 2.1 Microfluidic Phantom Fabrication

The microfluidic flow phantom was cast using PDMS (Sylgard 184™, Dow Corning) with 1.8 mg titanium dioxide (TiO_2_) added per gram of PDMS to generate a scattering background that mimics tissue optical properties at near infrared wavelengths (*μ*^*’*^_*s*_ = 8 cm^−1^).^14, 31^ The PDMS TiO_2_ mixture was vigorously mixed for 15 minutes to ensure even distribution of the TiO_2_ in the phantom. Centrifugation (3000 g relative centrifugal force, 10 minutes) was then performed to de-gas the sample and remove any TiO_2_ clumps.

A machined aluminum mold was used to cast the microfluidic as shown in Fig. 1a. The extrusion is 460 *μ*m × 460 *μ*m × 25 mm, which formed three walls of the channel. The square cross-section (1:1 aspect ratio) approximates the circular cross-section of vascular geometries *in vivo*. The PDMS TiO_2_ mixture was carefully poured on top of the mold within a polystyrene petri dish and placed in a vacuum chamber to remove air bubbles introduced during pouring. The PDMS was then cured at 70 °C for one hour, allowed to cool, and detached from the mold. The microchannel side of the phantom was oxygen plasma bonded to a glass slide to seal the channel. Inlet and outlet ports were attached using 23 gauge blunt tip needles fitted partially inside the phantom, which were epoxied in place for an air-tight seal. Tygon® microbore tubing (0.02^*’’*^ inner diameter, Cole-Parmer Instrument Company, LLC) was connected to the blunt tip ports as inlet and outlet tubing. The inlet tubing was connected to the flow control system using fluidic fittings with FEP tubing (IDEX Health and Science, LLC). The outlet tubing drained into a vial, with the tubing end fully submerged in liquid (see Supplementary Material S1). A photograph of the phantom is shown in Fig. 1b, where the channel is visible inside the white PDMS with the glass slide sealing the surface.

**Fig 1.**
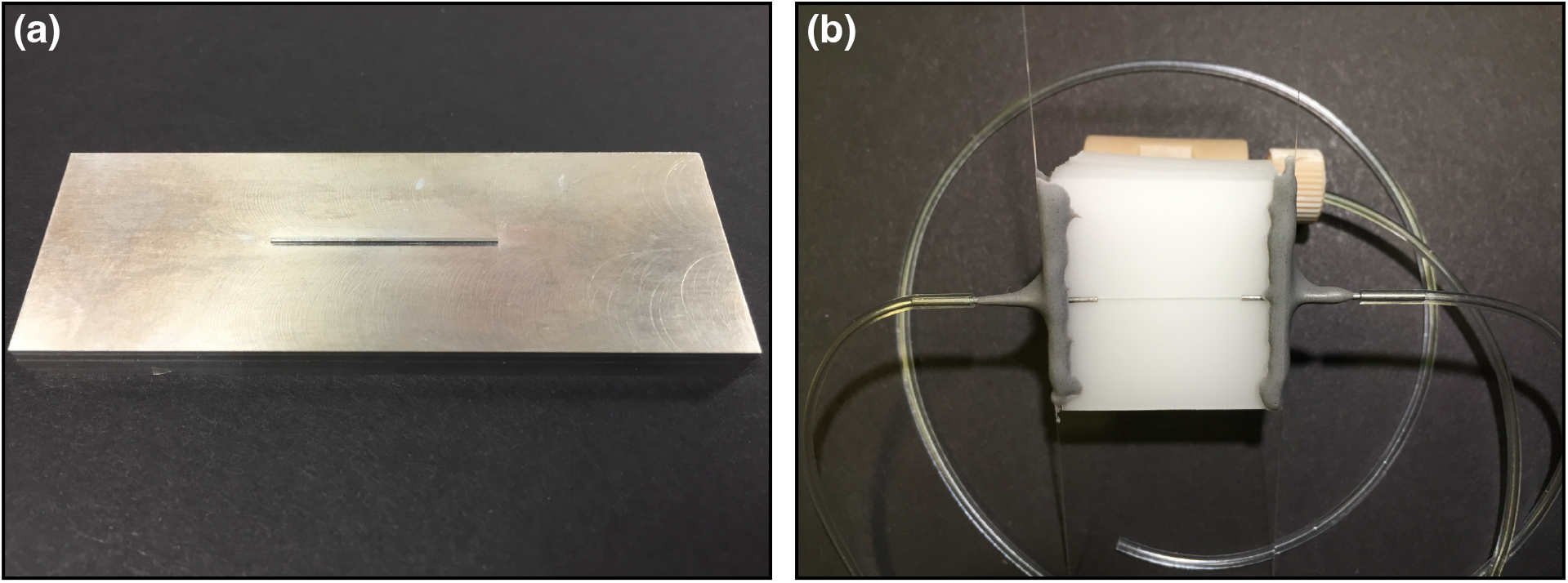
(a) Photograph of the machined aluminum mold with a 460 *μ*m × 460 *μ*m × 25 mm extrusion used to cast the PDMS during fabrication of the microfluidic flow phantom. (b) Photograph of the finished microfluidic flow phantom, showing PDMS bonded to a glass slide, with inlet and outlet ports secured using epoxy. Tubing was secured to the ports and connected with the rest of the system using fluidic fittings.

A colloidal mixture of suspended 1 *μ*m diameter polystyrene microspheres (10% w/w, 5100A, Thermo Fisher Scientific, Inc.) in ultra-filtered de-ionized (UFDI) water was used as a blood-mimicking solution. Based on Ref. 32, the reduced scattering coefficient (*μ*^*’*^_*s*_) of whole blood with hematocrit of 0.5 is approximately 20 cm^−1^. By using a 7% by volume mixture of the microsphere solution and UFDI water (e.g. 10 mL of blood mimicking solution = 0.7 mL of the 5100A solution + 9.3 mL UFDI water) we were able to achieve a calculated *μ*^*’*^_*s*_ of 19.8 cm^−1^. However, we note that the anisotropy of the microsphere solution is different than that of whole blood (*g* = 0.92 versus *g* = 0.98). The blood-mimicking solution was preferred for several reasons: cleaning and safety, ease of acquisition, and because the flow sensor was factory-calibrated for deionized water. Since the concentration of microspheres was very low (0.7% by weight in the final suspension), the water calibration of the flow sensor remained valid. The solution of microspheres was freshly mixed before each experiment to reduce particle aggregation. Once mixed, the solution was sonicated for 10 minutes to fully suspend the particles and then de-gassed under vacuum to eliminate air bubbles.

### 2.2 Microfluidic Flow System Instrumentation

To assess the performance of the syringe pump and pressure-regulated flow systems, and to quantify the uncertainty of each system, it is necessary to know the true flow rate of the blood-mimicking solution through the microfluidic channel. A thermal mass flow sensor (FLU-L, 0.03 – 1 mL/min, Fluigent, Inc.) was placed inline between the flow control system and the phantom inlet tubing, as shown in Fig. 2a. The sensor output was digitized and relayed to a computer via an accompanying hub (Flowboard, FLB, Fluigent, Inc.) where the MAESFLO software (Fluigent, Inc.) was used to monitor and record the absolute flow rate. All flow sensor measurements were downsampled from 10 Hz to 2 Hz using interpolation to match the timing of the optical measurements.

**Fig 2.**
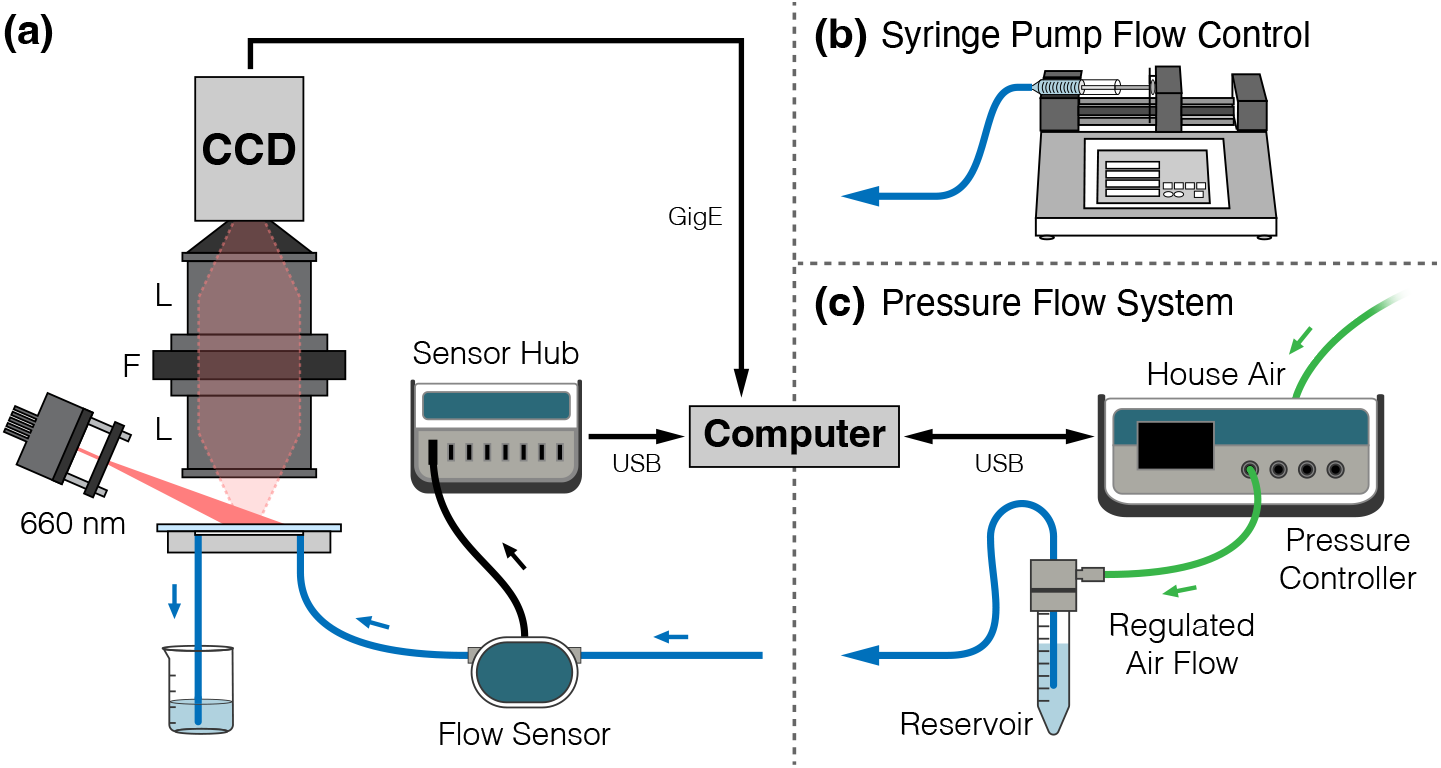
(a) LSCI system schematic for microfluidic flow assessment (L = Lens, F = Filter). A commercial flow sensor was placed in series with the inlet tubing for the microfluidic flow phantom to obtain real-time absolute flow measurements. Flow was programmatically controlled using either a (b) syringe pump or (c) pressure-regulated flow system. The software for the pressure flow system used a feedback loop between the flow sensor readings and the pressure controller to regulate air pressure.

The syringe pump system (74905-04, Cole-Parmer Instrument Company, LLC) was connected to the flow sensor (Fig. 2b) and a 10 mL plastic syringe (#302149, BD) was filled with the blood-mimicking solution described in Section 2.1. The syringe pump was programmed to run a predefined flow profile, using the 14.5 mm inner diameter of the syringe to ensure a correct flow rate. It should be noted that a 3 mL syringe was also evaluated, however its performance was worse than the 10 mL syringe and could not hold sufficient volume for all trials without being refilled.

The pressure-regulated flow system used a pressure controller (MFCS-EZ with 69 mbar channel, Fluigent, Inc.) connected to a house air line regulated at 500 mbar, as shown in Fig. 2c. This controller outputted regulated air flow (up to 69 mbar) to a 15 mL pressurized reservoir (Fluiwell-1C, Fluigent, Inc.) filled with the blood-mimicking solution. Prior to running a flow program, a calibration was performed that accounted for the resistance from the tubing, microfluidic channel, and hydrostatic pressure. The MAESFLO control software used feedback from the inline flow sensor to regulate the air pressure to achieve the desired flow rate.

### 2.3 Microfluidic Flow System Assessment

Two stepped flow profile experiments were conducted to assess each flow system using the commercial flow sensor. First, a 3-step flow profile was used to determine the uncertainty of each flow control system. The experiment lasted 240 seconds, beginning with 30 seconds of no flow (0 mm/s) followed by one full minute at each flow speed (2.4, 3.6, and 4.8 mm/s) before concluding with an additional 30 seconds of no flow. The experiment was repeated three times on each system. The second flow profile experiment consisted of a 13-step flow profile. The flow speed was once again varied from 2.4 to 4.8 mm/s, however in smaller 0.2 mm/s increments. As with the first experiment, the flow profile begins and ends with 30 seconds of no flow, with one full minute at each programmed flow speed (840 seconds total duration). The second experiment was also repeated three times on each flow control system.

To quantify and compare each flow system, we calculated the total combined uncertainty in accordance with National Institute of Standards and Technology (NIST) guidelines.^33, 34^ Three main sources of uncertainty were identified for inclusion in the uncertainty budget: repeatability, reproducibility, and measurement bias. Standard uncertainties were calculated for each flow step using 55 seconds of data, allowing for 5 seconds of transition from the previous flow state. All values were expressed as percentages relative to their respective means.

The uncertainty in repeatability, which is a measure of within-run variance, was defined as:

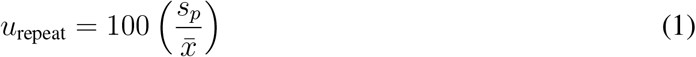

where 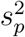 is the pooled variance across the three trials at each flow speed and 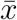 is the mean value of the measured flow speed across all trials.

The uncertainty in reproducibility, which is a measure of between-run variance, was defined as:

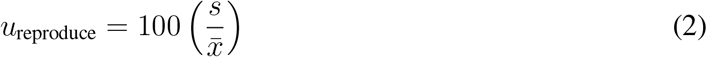

where *s*^2^ is the variance in the mean flow speed across all three trials and 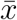 is the mean value of the measured flow speed across all trials.

Finally, the uncertainty in measurement bias, an estimate of flow accuracy, was defined as:

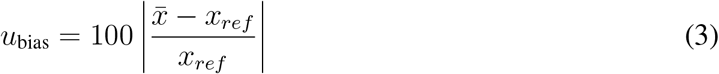

where 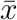 is the mean value of the measured flow speed across all trials and *x*_*ref*_ is the programmed flow speed (2.4, 3.6, or 4.8 mm/s) at each step.

The total combined standard uncertainty (*u*_*c*_) was then calculated using the root sum of squares of the individual uncertainty components and reported as the expanded uncertainty (*U* = *ku*_*c*_) with a *k* = 2 coverage factor corresponding to a level of confidence of approximately 95%.

### 2.4 LSCI Instrumentation

A schematic of the LSCI setup is shown in Fig. 2a. Measurements were performed using a 660 nm laser diode (HL6545MG, Thorlabs Inc.) mounted in a temperature-controlled housing (LDM21, Thorlabs Inc.) and expanded using an aspheric lens to obliquely illuminate the microfluidic. The laser diode controller (LDC205C, Thorlabs, Inc.) was set to 200 mA (∼120 mW) while the temperature controller (TED200C, Thorlabs Inc.) was set to 10.5 kΩ (23.8 °C). A pair of fixed focal length consumer camera lenses (AF Nikkor 50 mm *f* /1.8D, Nikon Corp.) were placed in tandem with infinity focus for 1:1 imaging. The aperture of the lower lens was set to *f* /11 in order to properly sample the speckle pattern^35^ while the upper lens was left at its maximum size of *f* /1.8. A longpass red filter (600 nm, R-60, Edmund Optics, Inc.) was placed between the two lenses to filter stray background light. The scattered light was imaged with a CCD camera (piA640-210gm, 648 × 488 pixels, Basler AG) controlled using custom software. The full sensor array was used, resulting in a field-of-view of 4.8 × 3.6 mm. The camera exposure time was set to 0.5 ms and allowed to run at full speed, resulting in an effective acquisition rate of ∼150 frames-per-second (fps).

### 2.5 LSCI Flow Measurements

To validate the flow sensor measurements, single-exposure LSCI was used to simultaneously image the microfluidic flow phantom during the 13-step flow experiments described in Section 2.3. Raw intensity images were converted to speckle contrast images using Eq. 4, where speckle contrast (*K*) is defined as the ratio of the standard deviation (*σ*_*s*_) to the mean intensity (*(I)*) in a sliding 7×7-pixel window centered at every pixel of the raw image.

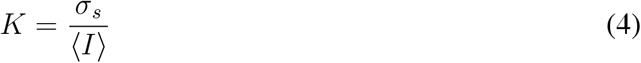

During post-processing, the average speckle contrast was extracted from each frame within a region centered in the microfluidic channel. This value was used to estimate the correlation time (*τ*_*c*_) of the speckle autocorrelation function, which is considered a more quantitative measure of flow.^36^ The averaged speckle contrast value at each timepoint was fitted for its corresponding *τ*_*c*_ using Eq. 5,

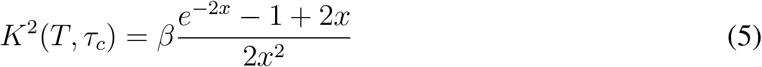

where *T* is the camera exposure time, *x* = *T/τ*_*c*_, and *β* is an instrumentation factor that accounts for speckle sampling, polarization, and coherence effects.^37^ For these measurements, it was assumed that *β* = 1, which exclusively allows for the computation of relative flow changes. The resulting *τ*_*c*_ estimates were resampled to 2 Hz using interpolation and smoothed temporally with a central moving average filter (*k* = 5).

Because *τ*_*c*_ is inversely related to the speed of moving scatterers in the sample,^38^ the inverse correlation time (*ICT* = 1*/τ*_*c*_) is commonly used for visualization. In order to compare the LSCI measurements with the flow sensor, the relative flow for each technique was calculated using the average value of the first step (2.4 mm/s) as the baseline. Because the flow sensor software used a separate clock than the LSCI acquisition, the datasets were synchronized during post-processing using the initial rise of the first flow step.

### 2.6 MESI Instrumentation

A schematic of the MESI setup is shown in Fig. 3. MESI was performed using a wavelength-stabilized 785 nm laser diode (LD785-SEV300, Thorlabs, Inc.) mounted in a temperature-controlled housing (TCLDM9, Thorlabs, Inc.) and collimated using an aspheric lens (C280TMD-B, Thorlabs, Inc.). The operating current on the laser diode controller (LDC205C, Thorlabs, Inc.) was fixed at 380 mA (300 mW) and the target temperature set to 25 °C on the temperature controller (TED200C, Thorlabs, Inc.). The collimated laser light was passed through a free-space optical isolator (Electro-Optics Technology, Inc.) to minimize back reflections that interfere with single frequency performance. Because the external volume holographic grating of the stabilized laser diode produces a dark spot in the far field, the laser was coupled into a single-mode fiber optic patch cable (P3-780A-FC-2, Thorlabs, Inc.) to obtain a Gaussian beam. The fiber output was re-collimated (F230APC-780, Thorlabs, Inc.), intensity modulated with an acousto-optic modulator (AOM, 3100-125; RF Driver 1110AF-AIFO-1.0, Gooch & Housego, Ltd.), and relayed to obliquely illuminate the microfluidic.

**Fig 3.**
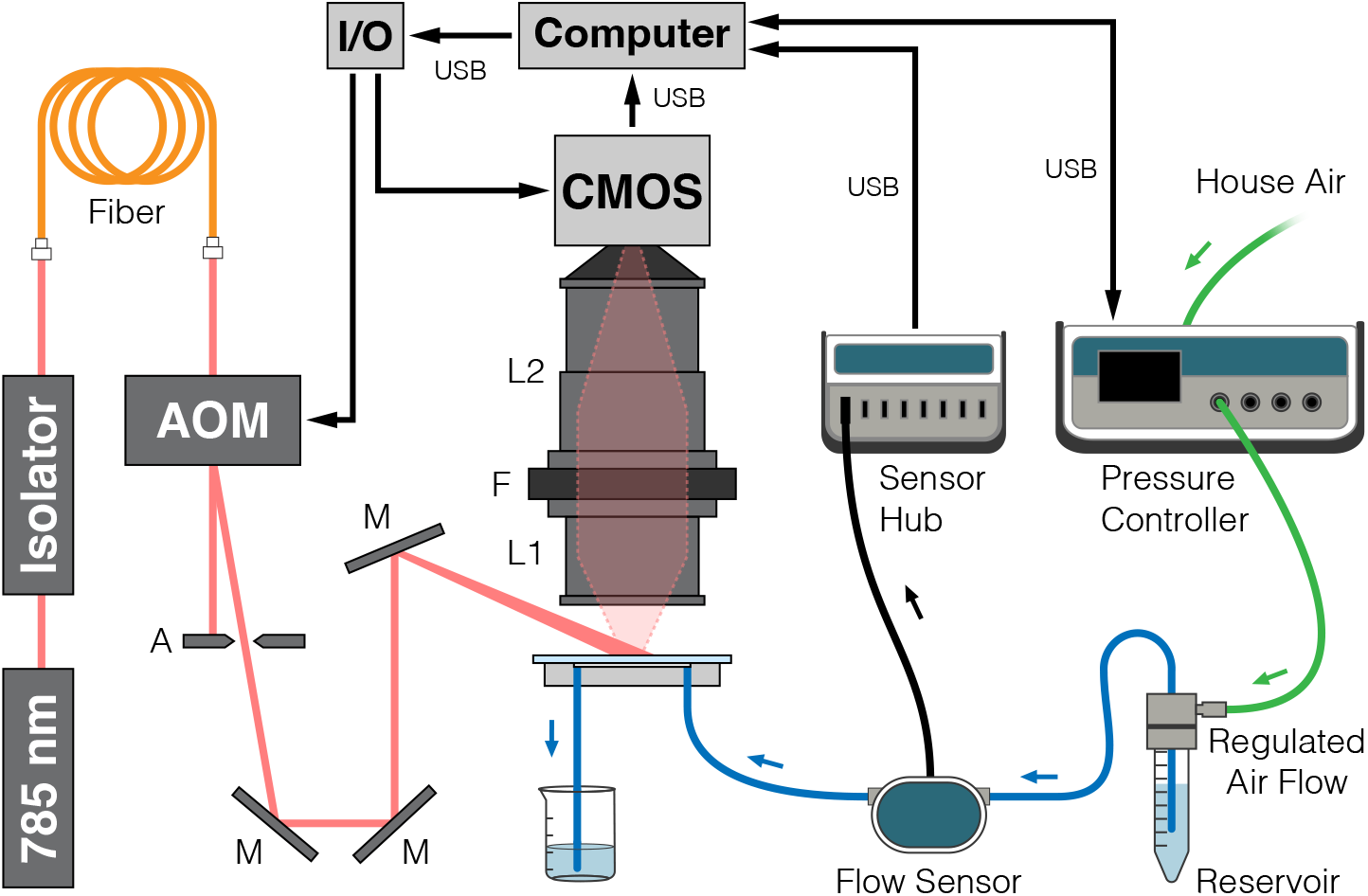
System schematic for microfluidic flow assessment of MESI and LSCI measurements using only the pressure-regulated flow system (A = Aperture, M = Mirror, L = Lens, F = Filter).

A pair of consumer camera lenses were used to image the scattered light (AF Nikkor 50 mm *f* /1.8D + AF Micro-Nikkor 105 mm *f* /2.8D, Nikon Corp.) with 2.1× magnification through a bandpass filter (785*±*31 nm, 87-773, Edmund Optics Inc.) to a CMOS camera (acA1920-155um, 1920 × 1200 pixels, Basler AG). The aperture of the lower lens was set to *f* /5.6 while the upper lens was left at its maximum size of *f* /2.8. Only a subset of the overall sensor array was used (1000 × 750 pixels), resulting in a field-of-view of 2.9 × 2.2 mm. The camera exposures were temporally synchronized with the modulated laser pulses. Fifteen different camera exposures ranging between 50 *μ*s and 80 ms were recorded for each complete MESI frame, resulting in an effective acquisition rate of approximately 2.5 fps. The total amount of light used to capture each exposure was held constant with the AOM in order to minimize the effects of shot noise.^14^ The entire acquisition was controlled using custom software written in C++ integrated with a multifunction I/O device (USB-6363, National Instruments Corp.) for the generation of camera exposure trigger signals and AOM modulation voltages.

### 2.7 MESI and LSCI Performance

A stepped flow profile experiment was conducted using the pressure-regulated flow system to assess the performance of both MESI and LSCI using the same optical hardware. The flow profile consisted of 19 steps spanning 1 – 10 mm/s in 0.5 mm/s increments. Each speed was maintained for two minutes with a total duration of 40 minutes, including brief periods of no flow (0 mm/s) at the beginning and end of the profile. MESI measurements were acquired continuously throughout the experiment at the maximum acquisition rate of the system (∼2.5 fps). Single-exposure (0.5 ms) LSCI frames were extracted from the MESI dataset *post hoc*. The initial rise of the first flow step was again used to synchronize the timing of the MESI/LSCI and flow sensor datasets. The experiment was repeated a total of four times across two days.

The raw intensity images were converted to speckle contrast images and the average value within the microfluidic channel extracted from each frame as described previously in Section 2.5. The average speckle contrast values from all fifteen exposure times were then fitted to the multi-exposure speckle visibility equation^14^ to obtain an estimate of *τ*_*c*_,

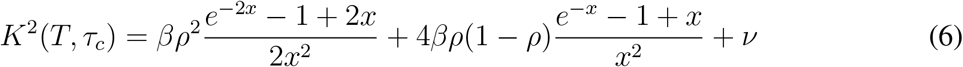

where *T* is the camera exposure time, *x* = *T/τ*_*c*_, *ρ* is the fraction of light that is dynamically scattered, *β* is a normalization factor that accounts for speckle sampling, and *?* represents exposure-independent instrument noise and nonergodic variances. *β* was held constant at all timepoints during an experiment by performing an initial fit to Eq. 6 using the median speckle contrast value at each exposure across the entire multi-exposure dataset. All fitting was performed with the Levenberg-Marquardt nonlinear least squares algorithm^39^ using a custom program written in MATLAB (R2021a, MathWorks, Inc.). The resulting *τ*_*c*_ estimates were resampled to 2 Hz using interpolation, smoothed temporally with a central moving average filter (*k* = 5), and inverted to obtain *ICT*.

In order to directly compare the MESI and single-exposure LSCI measurements with the flow sensor, the relative flow was calculated using the average value of the first step (1 mm/s) as the baseline. The average values for each flow step were calculated using 115 seconds of data, allowing for 5 seconds of transition from the previous step.

## 3 Results

### 3.1 Microfluidic Flow System Assessment

The comparison between the syringe pump and the pressure-regulated flow systems as measured with the inline flow sensor is shown in Fig. 4. These plots demonstrate that the pressure-regulated flow system outperforms the syringe pump by accurately and steadily matching the programmed flow speeds across both experiments. While both flow systems responded quickly to the desired flow changes during the large step size experiment (Fig. 4a), the syringe pump system suffered from accuracy and stability issues compared to the pressure-regulated flow system. Furthermore, the small step size experiment (Fig. 4b) demonstrated that the syringe pump system could not reliably reproduce the finer flow changes as evidenced by the loss of the stair-stepped flow pattern.

**Fig 4.**
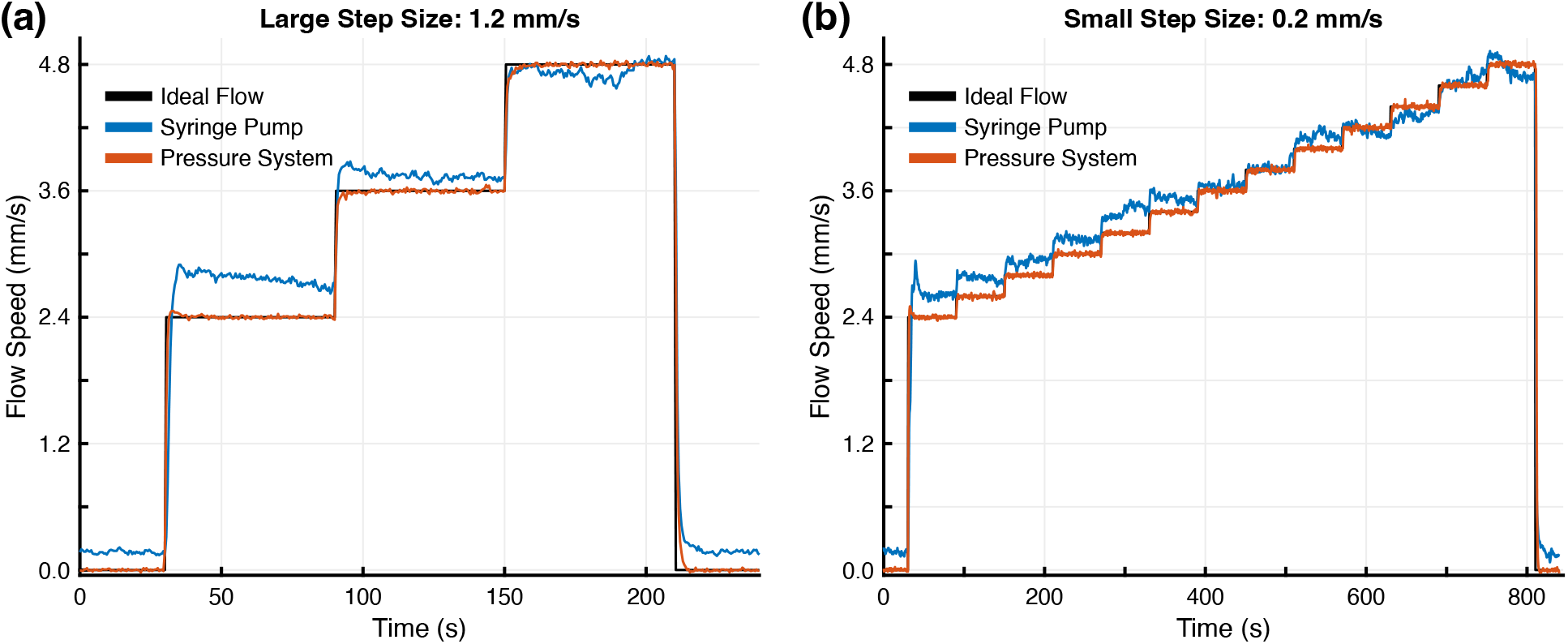
Flow sensor measurements for the syringe pump (blue) and pressure-regulated (red) flow systems compared to the ideal programmed flow (black) for one run of the (a) 3-step and (b) 13-step experiments. The microfluidic flow speed varied between 2.4 to 4.8 mm/s in 1.2 mm/s and 0.2 mm/s steps for the 3- and 13-step experiments, respectively.

Table 1 shows a comparison of the standard uncertainties for repeatability (*u*_repeat_), reproducibility (*u*_reproduce_), and bias (*u*_bias_) for each system at the flow speeds tested in the 3-step experiments. These results quantify the observations from Fig. 4 that the pressure-regulated flow system is more stable, repeatable, and accurate at all the tested flow speeds. The combined and expanded uncertainties for both systems are presented in Table 2. These results provide evidence of the superiority of the pressure-regulated flow system, particularly when attempting to produce small flow changes. With the syringe pump system, it would be impossible to reliably separate 20% changes in relative flow from flow system noise at the 2.4 mm/s flow speed. This magnitude of uncertainty in flow generation would make it difficult to assess an optical system’s ability to detect small changes in flow. This limitation is visually represented in Fig. 4b, where the syringe pump fails to resolve the finer 0.2 mm/s flow changes. Using the pressure-regulated flow system, experiments could be designed with relative flow changes of only 1.25% while remaining confident that the resulting optical system measurements were not caused by flow system noise. This level of certainty would also permit assessing improvements to optical system performance at this scale.

**Table 1.** Standard uncertainties for repeatability, reproducibility, and bias by flow speeds for the syringe pump and pressure-regulated flow systems.

| Flow Speed (mm/s) | $u_{\text{repeat}}$ (%) | | | $u_{\text{reproduce}}$ (%) | | | $u_{\text{bias}}$ (%) | | |
| --- | --- | --- | --- | --- | --- | --- | --- | --- | --- |
|  | 2.4 | 3.6 | 4.8 | 2.4 | 3.6 | 4.8 | 2.4 | 3.6 | 4.8 |
| **Syringe Pump** | 1.31 | 0.90 | 1.86 | 4.01 | 2.15 | 1.07 | 10.3 | 1.80 | 2.55 |
| **Pressure System** | 0.55 | 0.45 | 0.39 | 0.002 | 0.009 | 0.003 | 0.03 | 0.07 | 0.03 |

**Table 2.** Combined and expanded uncertainties (with *k* = 2 coverage) by flow speeds for the syringe pump and pressure-regulated flow systems.

| Flow Speed (mm/s) | $u_c$ (%) | | | $U$ (%) | | |
| --- | --- | --- | --- | --- | --- | --- |
|  | 2.4 | 3.6 | 4.8 | 2.4 | 3.6 | 4.8 |
| **Syringe Pump** | 11.1 | 2.94 | 3.34 | 22.2 | 5.89 | 6.68 |
| **Pressure System** | 0.56 | 0.45 | 0.39 | 1.11 | 0.91 | 0.77 |

### 3.2 LSCI Flow Measurements

The comparison between the syringe pump and the pressure-regulated flow systems as measured with the inline flow sensor and single-exposure LSCI using the system described in Section 2.4 is shown in Fig. 5. The underlying data for both plots are the same as in Fig. 4b but are normalized against the first flow step to permit comparison. The relative flow changes measured by LSCI closely track those of the flow sensor and corroborate the accuracy and stability issues seen with the syringe pump system. These results demonstrate that LSCI is capable of detecting similar changes as the commercial flow sensor when measuring relative flow changes up to double the baseline.

**Fig 5.**
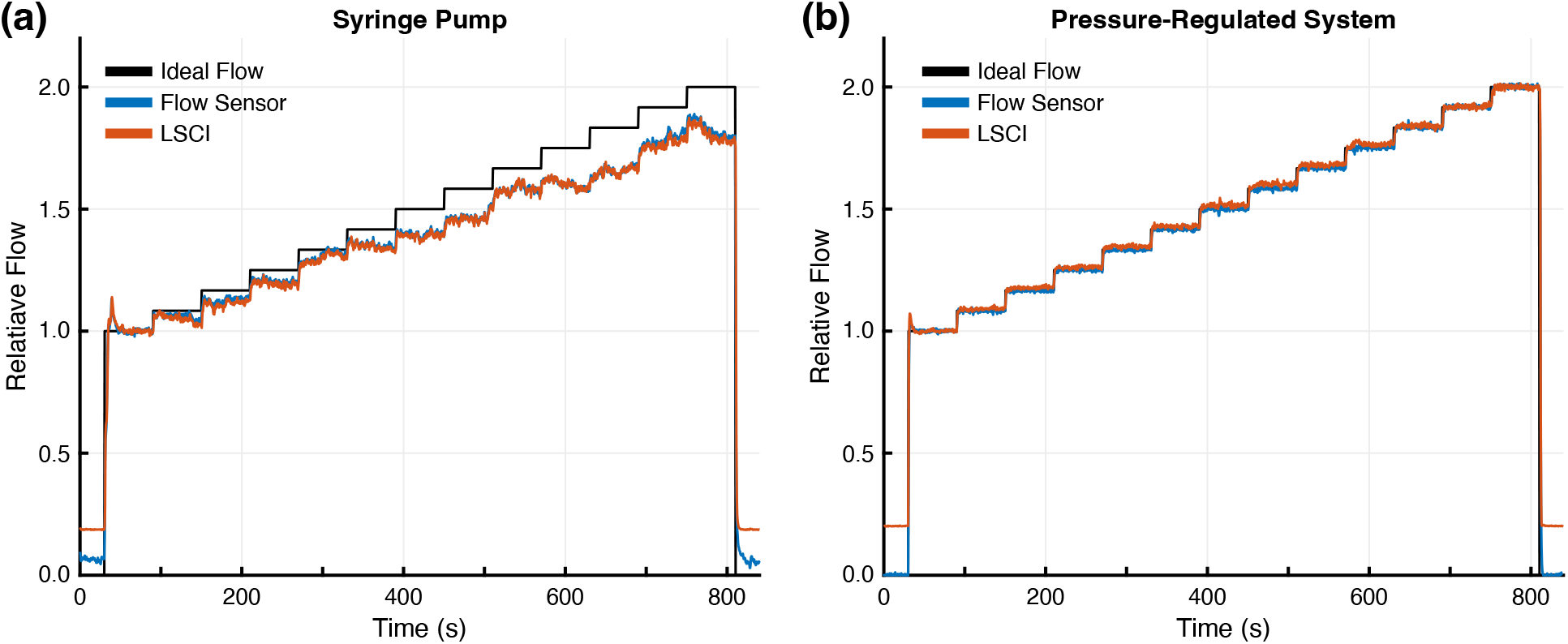
Relative flow measurements for the (a) syringe pump and (b) pressure-regulated flow systems as measured with the flow sensor (blue) and single-exposure LSCI (red). All measurements were normalized to the average of the first flow step (2.4 mm/s) and compared to the ideal programmed flow (black).

### 3.3 MESI and LSCI Performance

The flow measurement performance assessment using the pressure-regulated flow system is shown in Fig. 6. The relative flow measured by MESI and LSCI using the system described in Section 2.6 are plotted alongside the flow sensor measurements from a single trial in Fig. 6a. All relative flows were calculated using the average of the first flow step (1 mm/s) as the baseline. Data for each flow step were pooled and averaged across all four trials to assess the overall performance as seen in Fig. 6b. The coefficients of determination (*R*^2^) for this averaged MESI and LSCI data relative to the flow sensor were 0.988 and 0.839, respectively. The concordance correlation coefficients (*ρ*_*c*_), a metric designed to evaluate reproducibility,^40^ were 0.994 and 0.908, respectively. These results demonstrate both strong agreement between MESI and the flow sensor measurements and that MESI outperforms LSCI across large changes in relative flow. It should be noted that the apparent disparity in LSCI performance between Figs. 5b and 6 is primarily the result of different flow steps being used as the baseline in each set of experiments. While not an exact comparison, the LSCI measured flow at the 5 mm/s step is 1.94× the flow at the 2.5 mm/s step in Fig. 6, approximately mirroring the 2× change seen between the 2.4 and 4.8 mm/s steps in Fig. 5b.

**Fig 6.**
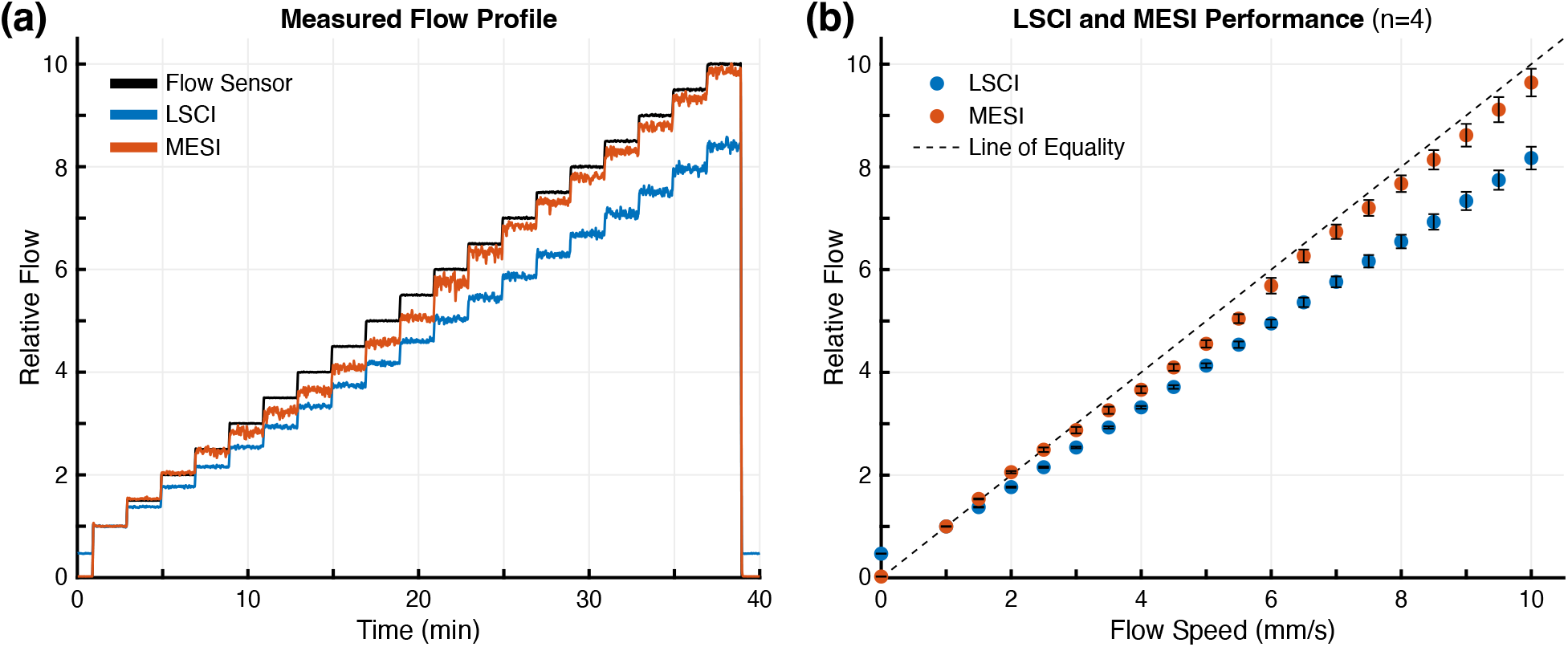
(a) Relative flow profiles for the flow sensor (black), LSCI (blue), and MESI (red) for one trial, all normalized to the average of the first flow step (1 mm/s). (b) Performance of LSCI and MESI flow estimates compared to measured microfluidic flow speed aggregated across all four trials (mean *±* sd). The dashed line denotes the line of equality indicative of a linear 1:1 relationship.

Fig. 7 summarizes the accuracy of the MESI and LSCI measurements at each flow step aggregated across all four trials. The absolute error for MESI was below 5% except for flow speeds between 3.5 to 6 mm/s, where it increased to almost 9%. In contrast, the LSCI error was over 8.2% for all flow speeds, increasing to over 17% for flow speeds greater than 4 mm/s. Aggregated across all tested flow speeds, the average absolute error for MESI was 4.6 *±* 2.6% and LSCI was 15.7 *±* 4.6%.

**Fig 7.**
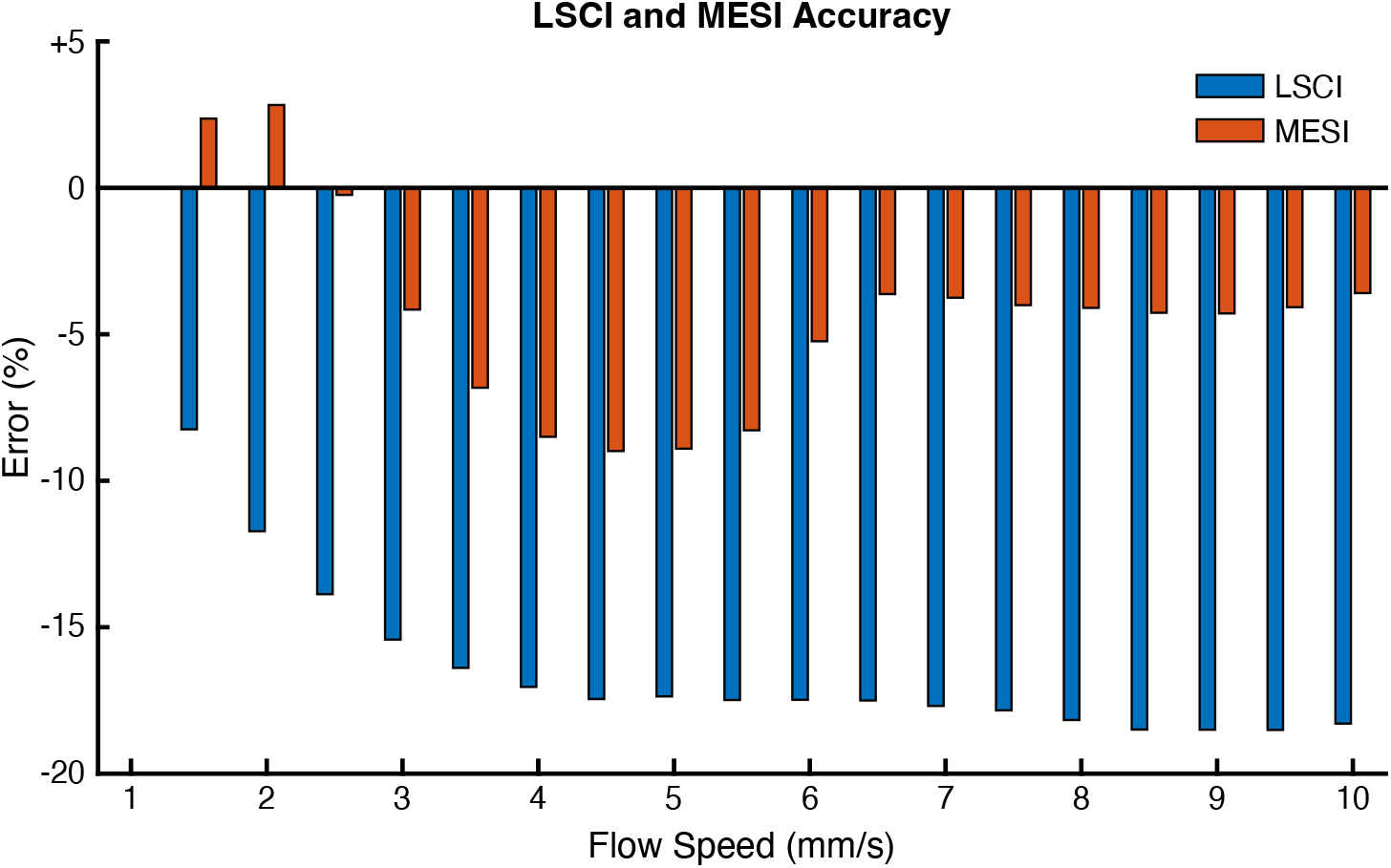
Percent error for LSCI (blue) and MESI (red) measurements at each flow step relative to the corresponding flow sensor measurements. Data aggregated by flow step and averaged across all four trials.

## 4 Discussion

Almost all microfluidic studies used for characterizing LSCI and MESI systems have been performed using syringe pumps. This study corroborates previous work^26, 28, 30^ that has shown syringe pumps do not produce stable flow outputs and demonstrates that single-exposure LSCI is sensitive enough to detect the resulting flow anomalies (Fig. 5). Furthermore, the close agreement between LSCI and the flow sensor demonstrates that these anomalies could be incorrectly interpreted as noise in the optical imaging system when they are actual flow variations. Compared to the syringe pump, the pressure-regulated flow system produced much more stable and reproducible flow outputs across all trials. Because microfluidic studies are commonly used to assess LSCI and MESI instrument performance, an unstable flow control system contributes additional noise to the measurements, making it more difficult to properly characterize a system. For example, the flow profiles in Fig. 4 and the uncertainty analysis in Table 2 show large errors and high variability within and across trials using the syringe pump. These instabilities in flow were caused entirely by the syringe pump, as independently validated using single-exposure LSCI in Fig. 5a. Such instability would increase the uncertainty in the true performance of any instrument being evaluated. The same analysis shows that a more reliable flow controller, such as the pressure-regulated flow system, produces minimal error and much improved reproducibility, therefore facilitating a more robust instrument assessment.

The measured noise levels quantified by *u*_repeat_ follow the specifications published by the manufacturers of both flow control systems. This indicates that the stability results shown are typical and representative of actual performance differences. The *u*_repeat_ for the syringe pump across all flow steps (up to 1.86%) was within the manufacturer’s tolerances, which state that instantaneous flow should be *±*2% from the mean value. The *u*_repeat_ for the pressure-regulated flow system (up to 0.55%) slightly exceeded the manufacturer’s specifications, which indicated that the best stability achievable with this system configuration was *±*0.5% from the mean value. Much larger differences arose from issues with reproducibility and accuracy, with the *u*_reproduce_ and *u*_bias_ of the syringe pump measuring several orders of magnitude higher than those of the pressure-regulated flow system. The syringe pump greatly exceeded the manufacturer’s tolerances for both reproducibility (*±*0.05%) and accuracy (*±*0.5%). Corresponding specifications were not available for the pressure-regulated flow system. While it is possible that using a glass syringe^28^ or sending the device for manufacturer recalibration could improve performance on the syringe pump, it is unlikely to ever achieve the stability and reliability seen with the pressure-regulated flow system.

The evaluation of MESI and single-exposure LSCI using the pressure-regulated flow system further demonstrates the superiority of the MESI technique.^16^ The MESI estimates of relative flow outperformed LSCI at all flow speeds and closely followed the flow sensor measurements. This included the periods with no flow at the beginning and end of each experiment, where LSCI still measured relative flow of almost 0.5× the baseline. As seen in Fig. 7, the *largest* absolute error for MESI was comparable to that of the *smallest* absolute error for LSCI. While MESI exhibited slightly more variability during individual flow profile steps (Fig. 6), this was likely caused by the fitting process required to calculate *τ*_*c*_. Numerical artifacts introduced by this nonlinear fitting could also be responsible for the increased error seen with MESI at flow speeds between 3.5 to 6 mm/s. It is also possible that discrepancies between MESI and the flow sensor could be attributed to a difference in flow between the microfluidic channel and the inlet line where the flow sensor is located. Such a difference could be caused by air leakage in the inlet or outlet port of the phantom, which would reduce the flow measured by the MESI system without impacting the upstream flow measured by the flow sensor.

The linearity of the results seen with both MESI and LSCI suggest that a calibration to absolute flow speeds might be feasible. However, any such calibration would be highly dependent upon experimental parameters, including the sample geometry (i.e. channel size), scatterer concentration, illumination configuration, and exposure time selection. Changes to any of these parameters would produce different *ICT* values, which would affect the correlation with the flow sensor and invalidate the calibration. It also remains to be seen whether calibrations performed in controlled microfluidic environments can reliably be translated to animal or human subjects since *in vivo* measurements sample a range of different vessels.^41, 42^

While the pressure-regulated flow system outperformed the syringe pump during testing, it has two major disadvantages: increased cost and difficulty of use. For the models used in this study, the pressure-regulated flow system was ∼3.5× more expensive than the syringe pump. Although there is variability in pricing for both types of systems depending on the vendor and specific features, pressure-regulated flow systems are consistently the more expensive. They are also considerably more difficult to use and maintain than a syringe pump, which simply requires filling a syringe with solution, securing it to the pump, and specifying the desired flow rate. The setup for a syringe pump can be performed quickly with minimal room for error. In contrast, the pressure-regulated flow system requires an extensive setup process and much greater care must be taken in order to achieve the stated performance of the device. For example, in order to maximize stability, tubing lengths must be selected to achieve a certain level of resistance in the system such that the desired flow range (0 – 60.9 *μ*L/min) spans the available range of the pressure controller (0 – 69 mbar). Furthermore, since fluidic fittings are required for connecting all system components, diagnosing instabilities caused by air leakage from an improperly secured fitting can become a difficult endeavor. The system also requires a calibration procedure before any flow program can be executed. While this helps ensure a high level of performance, it is an additional step during an already lengthy setup process. Overall, the pressure-regulated flow system was more tedious and had much greater potential for error. However, in order to accurately test instrument sensitivity to small flow changes, the pressure-regulated flow system was the only option with the necessary stability and reliability required to produce such flow changes.

## 5 Conclusion

This paper demonstrates that flow control system selection is critical for reliably generating flow for microfluidic studies. While ubiquitous in the LSCI community, syringe pumps suffer from poor stability and repeatability that limit their ability to properly characterize the flow sensitivity of optical imaging systems. We show that pressure-regulated flow systems should be used instead of syringe pumps when assessing the performance of MESI or LSCI with microfluidics. We determine that the uncertainty in flow generation with a pressure-regulated flow system is considerably less than that of a syringe pump system across a range of physiologically-relevant flow speeds. We use the pressure-regulated flow system to further demonstrate the superiority of MESI compared to traditional single-exposure LSCI. Using an inline flow sensor as reference, we show that MESI offers higher accuracy than LSCI across a wide range of flow speeds, with average errors of 4.6% and 15.7%, respectively. Given the large uncertainties associated with the syringe pump system, it would have been impossible to conclude that such measurements arose from an actual flow change rather than systematic error. Overall, these results indicate that highly accurate flow control is necessary for the reliable characterization of a MESI or LSCI system.

## Author Contributions

C.T.S., L.M.R., and P.K.M. constructed the imaging system, conceived the experiments, and acquired the data. C.T.S., A.S., L.M.R., P.K.M., and C.S. analyzed the results. C.T.S, A.S., and L.M.R. wrote the manuscript. A.K.D. supervised the project. All authors contributed to manuscript revisions.

## Acknowledgments

This study was supported by the National Institutes of Health (Nos. NS108484 and EB011556) and the UT Austin Portugal Program.

## Data, Materials, and Code Availability

The code and data that support the findings of this study are available from the corresponding author, A.K.D., upon reasonable request.

