## Supplementary Material for "Using pressure-driven flow systems to evaluate laser speckle contrast imaging"

### S1 Importance of submerging the microfluidic outlet line

An important methodological detail taken into account during all experiments was the submersion of the microfluidic outlet tubing in solution as depicted in the system schematics in Figs. 2 and 3. This strategy was adopted after visualizing dramatic changes in flow caused by the effluent dripping into the collection vial when the outlet line was elevated above the solution. The release of drops from the end of the tubing causes a large decrease in pressure, resulting in a flow decrease. This was observed with both flow control systems as shown in Fig. S1, indicating that this is an important experimental consideration for any microfluidic setup. These trials spanned ten minutes, with approximately five minutes of the effluent dripping into the collection vial before manually submerging the outlet line in the collected solution.

Some general trends were observed across these trials. First, the magnitude of the flow decrease was larger for the syringe pump compared to the pressure-regulated flow system. Because the pressure-regulated system has a higher compliance than the syringe pump, high frequency perturbations are dampened more by the pressure-regulated system. Second, the frequency of the droplets was very regular within individual trials. This suggests that drops must reach a critical volume to overcome the surface tension in order to release from the tubing. Using the absolute flow rate from the inline sensor and the time between drops to estimate the mean volume, a standard deviation of $<0.5$ $\mu$L in drop volume was observed within each trial. Finally, the magnitude of the flow decrease was larger for the slower flow (2.4 mm/s) than the faster flow (4.8 mm/s) across all trials. This was likely caused by the larger pressure differential generated at slower flows, since the pressure driving the flow was lower for the slower flow rate. For example, with the pressure-regulated system, the drops led to a 1.4 mbar change at the 2.4 mm/s speed (28.3 mbar average) and a 1.1 mbar change at the 4.8 mm/s speed (47.4 mbar average). The larger relative pressure differential (4.95% vs. 2.3%) produced a larger change in flow, which corresponds to the higher magnitude flow decreases seen at the slower flow rate. Overall, these results demonstrate that users should be aware of the positioning of the microfluidic outlet line during experiments and that submersion of the outlet tubing is critical to eliminate periodic flow perturbations caused by dripping.

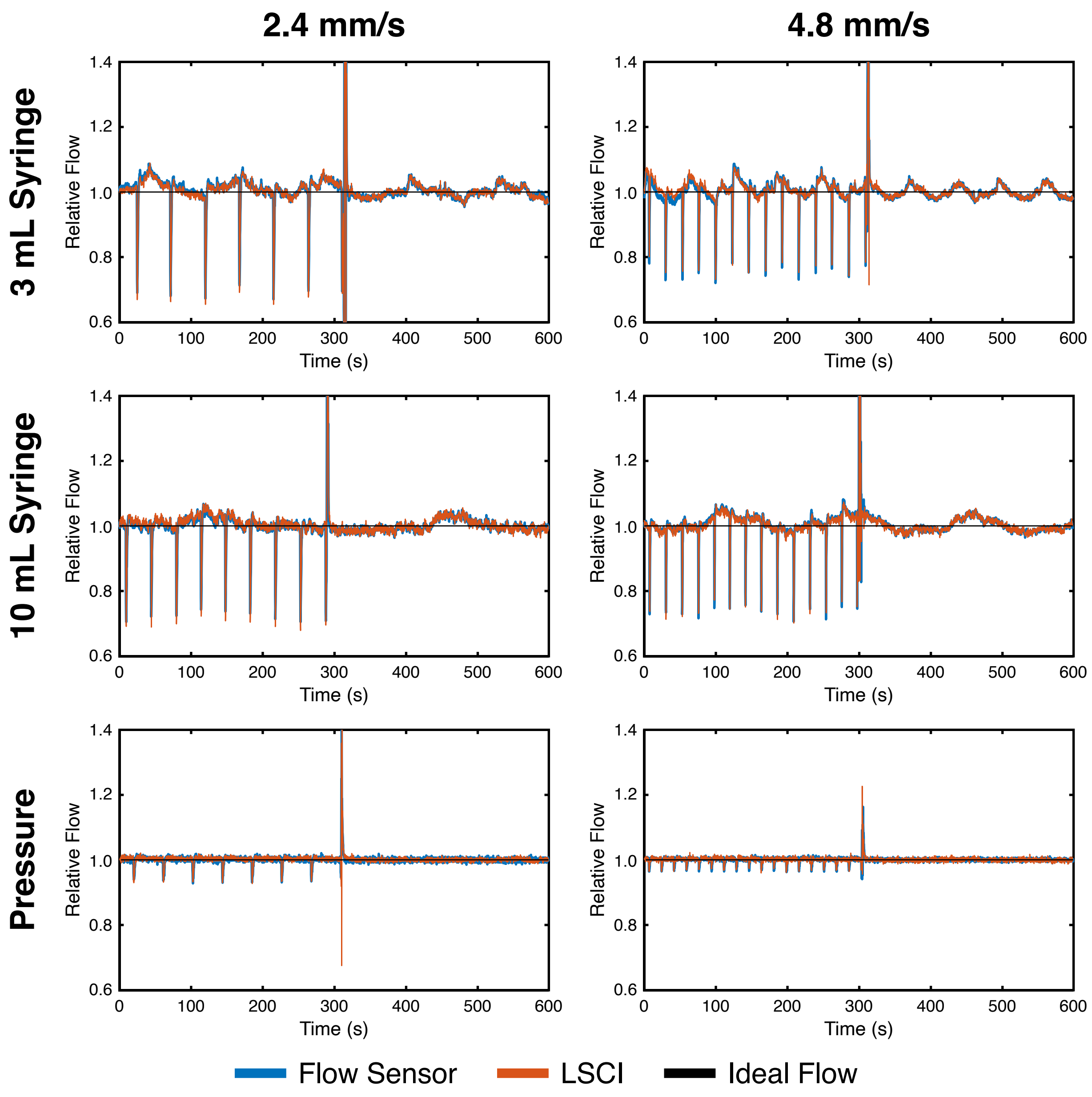


Figure S1: Impact of submerging the microfluidic device outlet tubing for both flow control systems at 2.4 and 4.8 mm/s as measured with the inline flow sensor (blue) and single-exposure LSCI (red). These trials show approximately five minutes of effluent dripping into the collection vial, as evidenced by the periodic flow decreases, followed by five minutes of operation with the outlet tubing submerged in the collected solution. The large spike in relative flow around $t = 300$ seconds corresponds to the outlet line being manually submerged in the solution. All measurements were normalized to the average flow during the submerged segment of each trial and compared to the ideal constant flow (black).
